# Antagonistic regulation of *Drosophila* mitochondrial uncoupling protein *UCP4b* by cold and BMP signaling

**DOI:** 10.1101/2022.01.27.477603

**Authors:** Nirmalya Chatterjee, Wei Song, Phillip A. Dumesic, Bruce Spiegelman, Norbert Perrimon

**Affiliations:** Department of Genetics, Blavatnik Institute, Harvard Medical School, Boston, MA 02115, USA; Dana-Farber Cancer Institute, Boston, MA 02115, USA; Department of Cell Biology, Blavatnik Institute, Harvard Medical School, Boston, MA 02115, USA; Howard Hughes Medical Institute, Boston, MA 02115, USA; Frontier Science Center for Immunology and Metabolism, Medical Research Institute, Wuhan University, Wuhan, Hubei 430071, China

## Abstract

Regulation of energy metabolism and response to cold are intimately linked in mammals. Central to these two processes are the mitochondrial uncoupling proteins (UCPs), which by promoting proton leakage across the inner mitochondrial membrane lead to the generation of heat instead of ATP synthesis. In addition to heat generation, UCPs also influence energy storage and can protect against obesity and diabetes. Cold-blooded animals like flies also contain UCPs that protect from cold, however their regulations are poorly understood. We find that *Drosophila UCP4b* is induced by cold in a cell-intrinsic manner and protects against cold and obesity in fly models. Mechanistically, cold regulates *UCP4b* expression through calcium signaling and Spargel (Srl), the *Drosophila* ortholog of mammalian PGC1α. To the opposite, MAD, acting downstream of the BMP branch of the TGFβ signaling pathway, represses *UCP4b* expression independently of cold. Interestingly, the two mechanisms of *UCP4b* regulation are integrated as MAD binding to the *UCP4b* promoter is displaced by cold in a *Srl-*dependent manner. We discuss the similarities between the regulation of mammalian and *Drosophila* UCPs.

**Significance:** Mitochondrial uncoupling proteins (UCPs) that uncouple the mitochondrial respiration from ATP synthesis regulate energy metabolism, non-shivering thermogenesis, and redox balance in vertebrates and invertebrates. However, their regulation in *Drosophila* is poorly understood. We found that *Drosophila* uncoupling protein UCP4b is induced by cold in a cell-autonomous fashion. Conversely, MAD, acting downstream of BMP signaling, inhibits *UCP4b* expression. MAD is displaced from the upstream regions of the *UCP4b* gene by cold. UCP4b protects *Drosophila* against cold and diet-induced obesity. The regulation of UCP4b by cold and BMP signaling is reminiscent of the regulation of mammalian uncoupling protein UCP1. Altogether, we discovered an important regulator of *Drosophila* energy metabolism which is controlled by regulatory processes that are similar between *Drosophila* and mammals.

## Introduction

Mitochondrial uncoupling proteins (UCPs), a family of transmembrane proton carriers, can cause proton leakage across the inner mitochondrial membrane and dissipate the proton gradient generated by the electron transport chain in the form of heat. This dissociates mitochondrial respiration from the function of ATP synthase that uses the proton gradient to generate ATP (1). UCPs play important roles in non-shivering thermogenesis (NST), a method of conversion of metabolic energy into heat without skeletal muscle contraction. In particular, mammalian mitochondrial uncoupling protein UCP1 (thermogenin), which is induced in brown adipose tissue (BAT) in response to cold, plays a pivotal role in NST (2). In addition to thermoregulation, mammalian UCPs reduce mitochondrial reactive oxygen species (ROS) generation by reducing the proton gradient across the inner mitochondrial membrane. For example, UCP2 expression is reduced in macrophages with a concomitant increase in ROS that is required to prevent infection (3), whereas UCP3 protects the heart against ischemia-reperfusion (IR) injury, which causes oxidative stress (4). Similarly, reduction in the expression of UCP4 and UCP45 in the dopaminergic neurons is implicated in oxidative stress in the DJ-1 (PARK7) mouse model of Parkinson’s disease (5).

The regulation of the expression of mammalian *Ucp1* by cold is fairly well characterized. Norepinephrine released in response to cold exposure triggers β-adrenergic signaling in brown adipose tissue (BAT), leading to the induction of *Ucp1* expression in BAT (2). In addition, the transcriptional co-activator *Ppargc1a* is induced in the mammalian BAT by cold-triggered β-adrenergic signaling and is involved in cold-mediated *Ucp1* induction in the BAT (6).The absence of PGC1α in brown adipocytes impairs the activation of the thermogenic gene expression program in response to cold (7). Interestingly, cold exposure can also increase thermogenic gene expression including *Ucp1* and *Ppargc1a* in white and beige adipocytes in a cell-autonomous fashion (8). This yet uncharacterized mechanism is different from the cold-mediated *Ucp1* induction in BAT (8). On the other hand, TGFβ/BMP signaling suppresses mammalian *Ucp1* expression. Smad3, acting downstream of TGFβ signaling, represses *Ucp1* expression in mammalian white adipose tissue (WAT) (9). Moreover, the induction of *Ucp1* under *Smad3* loss-of-function condition is also accompanied by an induction of *Ppargc1a*. Interestingly, *Smad3* deficiency prevents diet-induced obesity in mice (9). Similarly, BMP4, which promotes a brown to white-like adipocyte shift, inhibits *Ucp1* expression (10). Interestingly, BMP4 also suppresses *Ppargc1a* expression (10).

Insects, being cold-blooded animals employ processes such as cold avoidance, accumulation of cryoprotectants in the hemolymph, and changes in the membrane composition in response to cold (11–13). However, recent studies have indicated that mechanisms similar to mammalian non-shivering thermogenesis are also involved in cold tolerance in insects. There are four mitochondrial UCPs in *Drosophila*, namely UCP4a, UCP4b, UCP4c, and UCP5 (14). UCP4c promotes mitochondrial uncoupling and is required for the development of larvae when raised at cold temperatures (15). In addition, the expression of UCP4b and UCP4c is suppressed by the circadian rhythm master-regulator gene *period* (*per*). *per* mutants show a high metabolic rate due to mitochondrial uncoupling and are more tolerant to cold. Interestingly, disrupted expression of UCP4b and UCP4c reverts the increase in mitochondrial respiration and enhanced cold tolerance in *per* mutant flies (16). Consistent with redox-regulatory effects of mammalian UCPs, *Drosophila* UCPs are also implicated in redox regulation. The periodic expression of *UCP4b* and *UCP4c* orchestrated by *per* reduces the ROS load in *Drosophila* intestinal stem cells (ISCs) and maintains their proliferative homeostasis (16). Moreover, *UCP4a* overexpression in the pink1/parkin *Drosophila* models of Parkinson’s disease reduce ROS levels and protect against oxidative stress (17). Thus, like mammalian UCPs, *Drosophila* UCPs are also involved in both cold tolerance and redox regulation indicating a conservation of physiological functions of UCPs between *Drosophila* and mammals. In addition to *Drosophila* UCPs, *Spargel (Srl)*, similar to its mammalian ortholog PGC1α, has also been implicated in cold tolerance as a mutation in *Srl* renders *Drosophila* susceptible to cold (18).

While UCP-mediated non-shivering thermogenesis mechanism is functional in *Drosophila*, it is not known how *Drosophila* UCPs are regulated. Here, we report that *Drosophila UCP4b* is induced in a cell-autonomous fashion by cold and is suppressed by BMP signaling. Moreover, we found that Srl plays a central role in the regulation of *UCP4b* by both cold and BMP signaling. Our study demonstrates that the role of some proteins and signaling pathways involved in the regulation of mitochondrial bioenergetics is generally evolutionarily conserved between *Drosophila* and mammals.

## Results

### *UCP4b* is induced by cold in a cell-autonomous fashion

To determine if *Drosophila* UCPs, like mammalian UCP1, can be induced by cold, we tested the expression of *UCPs* after cold exposure of different durations. Strikingly, only *UCP4b* is induced in response to cold exposure in adult flies (Fig 1A), and overexpression of *UCP4b* protects against cold (Fig 1B), whereas *UCP4b* knockdown makes them more susceptible to cold (Fig 1C). Similarly, *UCP4b* is significantly induced in *Drosophila* larvae in response to cold exposure (Fig 1D), and only *UCP4b* is induced by cold in multiple larval tissues such as the adipose tissue (the fat body), the skeletal muscle (the body wall), the gut, and the brain (Fig 1E, Fig S1, S2). Interestingly, *UCP4b* is also induced by cold in S2R+ and Kc *Drosophila* cells (Fig 1F, Fig S3), indicating that these cells can directly sense cold and can induce *UCP4b* independently of any neuronal input. Thus, cold induces *UCP4b* transcription in adult flies, multiple larval tissues, and in cultured *Drosophila* cells. Both mammalian and *Drosophila* UCPs have been shown to cause mitochondrial depolarization (2, 15, 16). To test the effect of UCP4b on mitochondrial potential, we generated a stable S2R+ cell line expressing *UCP4b* under a metal-inducible promoter. These cells were stained with TMRM which accumulates in mitochondria with higher membrane potential. Copper sulfate induced *UCP4b* overexpression reduced the TMRM signal indicating mitochondrial depolarization (Fig 1G). Interestingly, genetic and pharmacological upregulation of UCP1 reduces obesity in mammals (19). Similarly, ubiquitous expression of *UCP4b* in adults protects against high sugar-diet (HSD) induced increase in TAG levels (Fig. 1H).

**Figure 1.**
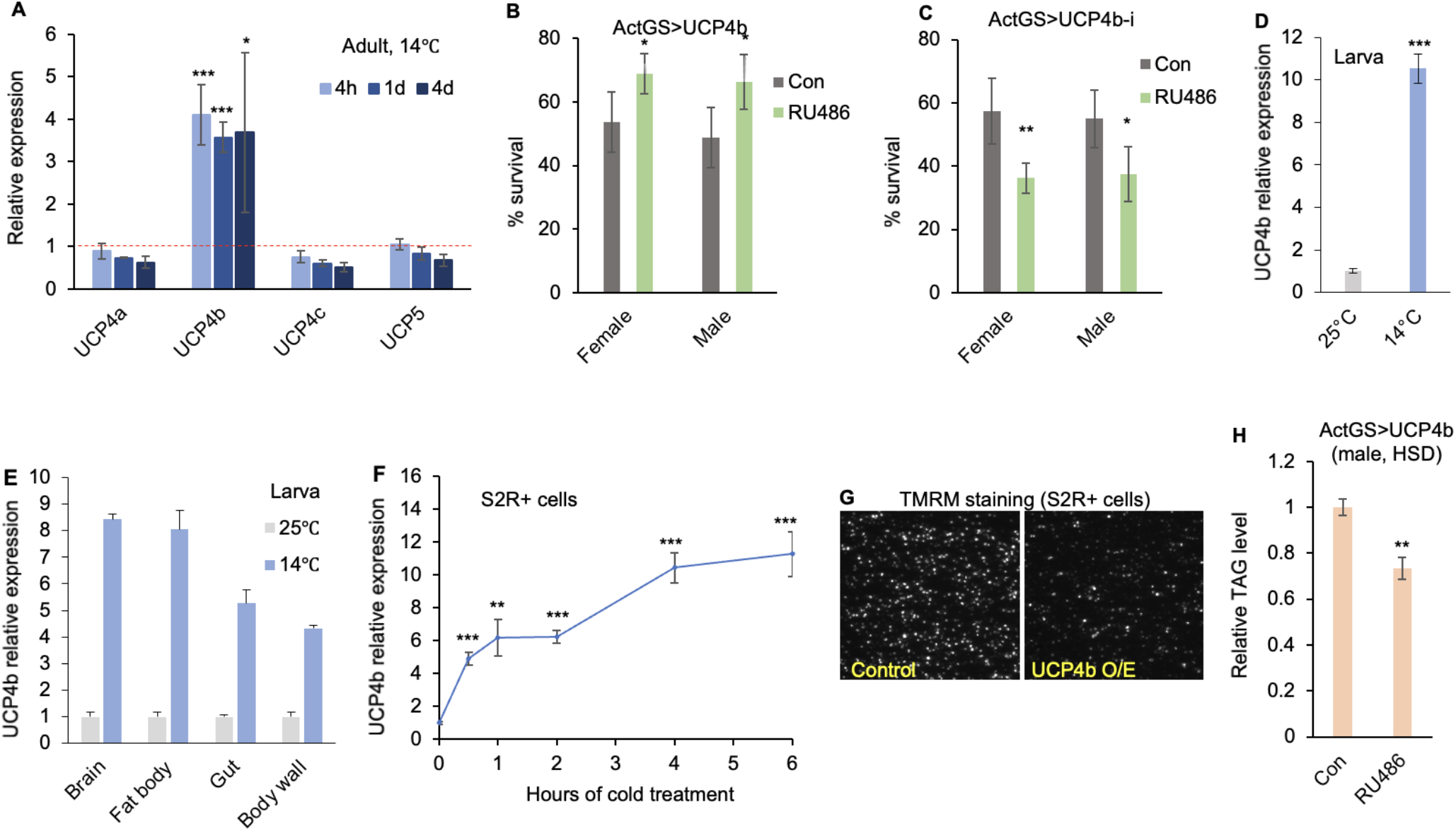
*UCP4b* is induced by cold in a cell-autonomous fashion and protects against cold. (A) 1-week old adult flies were exposed to cold (14°C) for different durations (4 hours, 1 day, and 4 days) followed by whole-body RNA extraction and qPCR (n=3). Transcript levels, normalized to α-tubulin, are expressed relative to their levels at 25**°**C. Error bars= SD; two-tailed unpaired t-test is performed to measure statistical significance [*(P<0.05); **(P<0.01); ***(P<0.001)]. (B) *UCP4b* was overexpressed with the ubiquitous Act-GeneSwitch (ActGS) driver that is inducible by the drug RU486 (ethanol is the vehicle control), and the cold tolerance of flies treated with RU486 and ethanol control was compared. (C) *UCP4b* was knocked down in adult flies under the control of ActGS. Flies were exposed to 4°C for 6 days and then were brought back to 25°C. The survivors were scored after 4 hours. Error bars= SD; two-tailed unpaired t-test is performed to measure statistical significance. (D) 3^rd^ instar *Drosophila* larvae were exposed to cold (14°C) for 6 hours followed by whole-body RNA extraction and qPCR (n=3). (E) 3^rd^ instar *Drosophila* larvae were exposed to cold (14°C) for 4 hours. Then the fat body, brain, gut, and body wall were dissected followed by RNA extraction and qPCR. (F) *Drosophila* S2R+ cells were exposed to cold (4°C) at different time points. Then RNA was extracted and qPCR was performed (n=3). Transcript levels are normalized to α-tubulin. (G) S2R+ cells stably transfected with MT-UCP4b plasmid were treated with copper sulfate or mock-treated for 48 hours. Then they were treated with oligomycin A, stained with TMRM, and images were collected. Representative images are shown here. (H) *ActGS>UCP4b* adult flies put on high-sugar food (15% w/v added sucrose) containing either RU486 or ethanol control for 10 days. Whole-body triglyceride (TAG) levels were measured in male flies (n=3, 8 adult males per replicate). The TAG levels are normalized to the number of flies. TAG levels are expressed relative to the values in flies on control food.

### Calcium signaling and Spargel (Srl) are involved in the cold-mediated *UCP4b* induction

Calcium signaling has been implicated in brain-independent *ex vivo* cold-sensing and rapid cold hardening (RCH) in insect tissues (20). Thus, we tested the role of calcium signaling in the cold-mediated induction *UCP4b* by monitoring the activity of the genetically encoded calcium indicator GCaMP in dissected larval fat body. *ex vivo* application of cold to larval adipose tissue activates calcium signaling (Fig 2A; Supplementary movie S1) and induces *UCP4b* (Fig S4). Moreover, pre-treatment with the chelator BAPTA-AM attenuates cold-mediated *UCP4b* induction in S2R+ cells indicating that calcium signaling plays a role in cold-mediated induction of *UCP4b* (Fig 2B). Cold-mediated *UCP4b* induction is not affected by pre-treatment with cycloheximide indicating that no intermediate protein synthesis is required for the process (Fig 2C). Moreover, *UCP4b* induction is completely attenuated by pre-treatment with actinomycin D (Fig 2D), indicating that cold-mediated increase in *UCP4b* transcript levels is principally caused by enhanced transcription of the *UCP4b* gene. Interestingly, knockdown of *Srl*, the ortholog of mammalian transcriptional co-activator *Ppargc1a*, strongly suppressed cold-mediated *UCP4b* induction in S2R+ cells (Fig 2E). However, *Srl* is not induced in response to cold either in cultured S2R+ cells (Fig 2F) nor in adult flies (Fig S5). Interestingly, ubiquitous *Srl* overexpression in larvae is sufficient to induce *UCP4b* in whole larvae (Fig 2G). Moreover, consistent with a previous study (18), *Srl* knockdown renders flies more susceptible to cold (Fig 2H). Thus, both calcium signaling and Srl are required for cold-mediated *UCP4b* induction.

**Figure 2.**
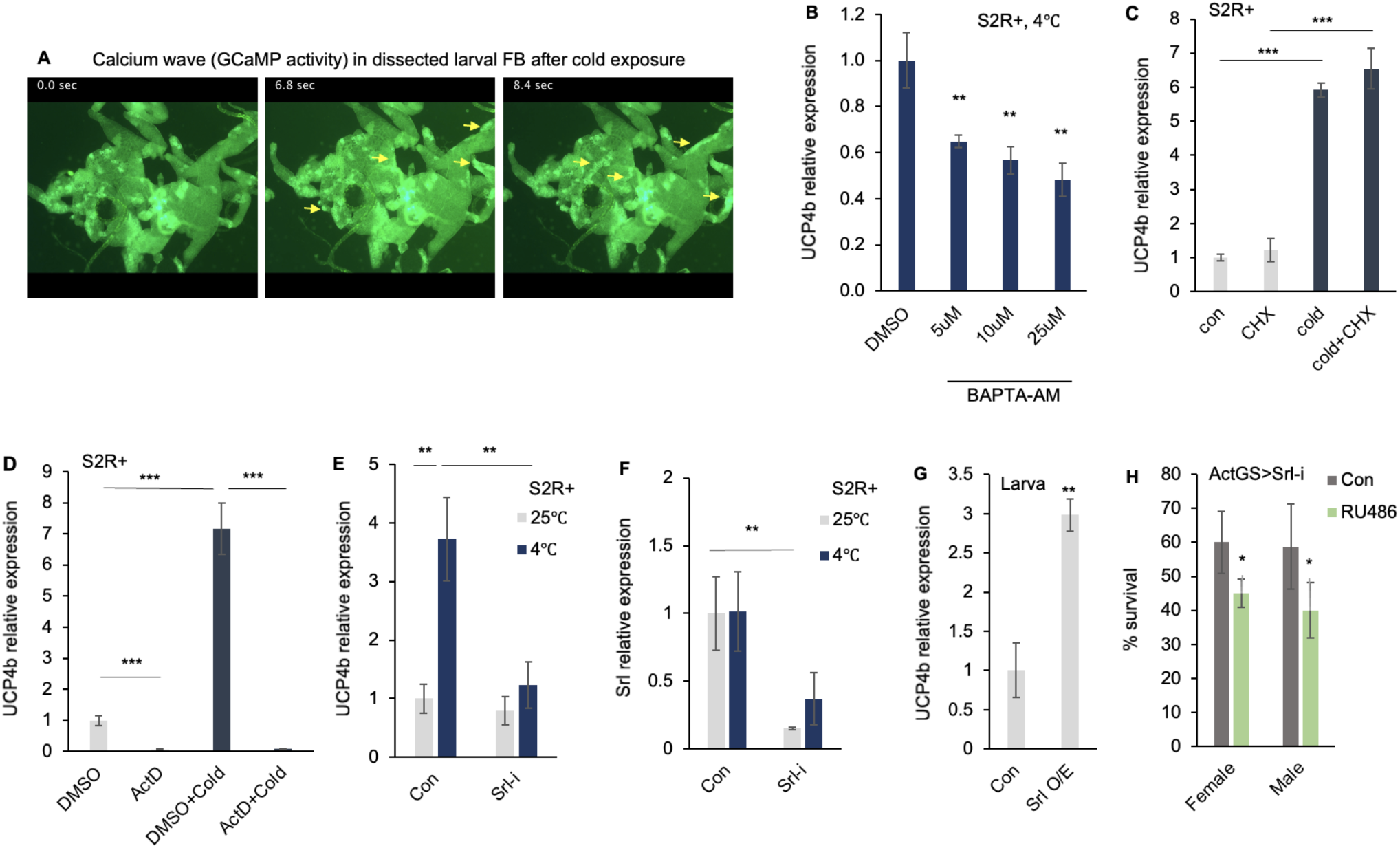
Calcium signaling and *Spargel (Srl)* are required for cold-mediated *UCP4b* induction. (A) The fat body was dissected in Schneider media at room temperature from 3^rd^ instar *Drosophila* larvae expressing calcium-sensitive GCaMP reporter under fat body-specific FB-Gal4 driver. Then cold (4°C) Schneider media was added and the changes in calcium levels were recorded under a microscope as a movie. The rise in GCaMP activity was initiated in multiple parts of the fat body and propagated in the form of waves to other parts. The images represent the temporal progression of the waves (see Supplementary Movie S1). The yellow arrows indicate locations with differential GCaMP activity in different timeframes (B) S2R+ cells were pre-incubated with calcium chelator BAPTA-AM before they were exposed to cold treatment (4°C for 1 hour). Then RNA was extracted and qPCR was performed; (n=3, normalized to α-tubulin). The relative *UCP4b* expression is compared to control pre-incubated with DMSO. (C) *Drosophila* S2R+ cells were pre-treated with 10 ug/mL cycloheximide (CHX) or mock-treated for 4 hours and then exposed to cold (4°C) for 2 hours. Then RNA was extracted and qPCR was performed. (D) *Drosophila* S2R+ cells were pre-treated with 10 ug/mL actinomycin D (ActD) or mock-treated for 4 hours and then exposed to cold (4°C) for 2 hours. Then RNA was extracted and qPCR was performed (n=3). Transcript levels are normalized to α-tubulin. (E, F) *Srl* was knocked down in S2R+ cells by dsRNA treatment, then the cells were exposed to cold (4°C for 1 hour). RNA was extracted and *UCPb* and *Srl* transcript levels were measured by qPCR (n=3, normalized to α-tubulin). (G) *tubGal4, tubGal80*^*ts*^; *UAS-Srl larvae* were exposed to a permissible temperature (29°C) for 2 days to allow UAS transgene expression and then *UCP4b* expression was compared with similarly treated *tubGal4, tubGal80*^*ts*^ larvae. (H) *Srl* was knocked down in adult flies with ActGS driver, and survivors were counted after exposure to 4°C following the protocol described in Fig 1C.

### *UCP4b* is suppressed by BMP signaling

MAD and its interacting partner MED, which is the ortholog of mammalian SMAD4, have been shown to bind to the upstream regions of *UCP4b* (21). These prompted us to explore the role of BMP signaling in the regulation of *UCP4b*. Interestingly, knockdown of *MAD* in S2R+ cells as well as knockdown of other components of BMP signaling such as *MED*, Type I receptors *Tkv* and *Sax*, Type II receptor *Put*, and *Gbb*, the only BMP ligand expressed in S2R+ cells, induced *UCP4b* expression under regular culture conditions at 25°C (Fig 3A). This indicates that BMP signaling suppresses *UCP4b* under basal conditions in both *Drosophila* larval fat body and in cultured cells. Moreover, the application of LDN212854, which has been reported to inhibit mammalian BMP signaling (22), leads to rapid induction of *UCP4b* expression in S2R+ cells (Fig 3B). This LDN212854-mediated *UCP4b* induction is not affected by a pre-treatment with cycloheximide (CHX), indicating that no intermediate protein synthesis is required for the process (Fig 3C). In addition, *UCP4b* was induced in adult flies fed a diet containing LDN212854 (Fig. 3D). On the other hand, ubiquitous overexpression of *MAD* in larvae suppressed cold-induced *dUCP4b* expression (Fig 3E). Similarly, *MAD* overexpression in the larval fat body suppressed *UCP4b* expression in the fat body (Fig 3F). Similar to the cold-mediated *UCP4b* induction, the induction of *UCP4b* in response to BMP signaling inhibition is dependent on Srl but *Srl* is not induced under these conditions (Fig 3G, H), indicating that Srl plays a central role in the regulation of *UCP4b* expression by both cold and BMP signaling.

**Figure 3.**
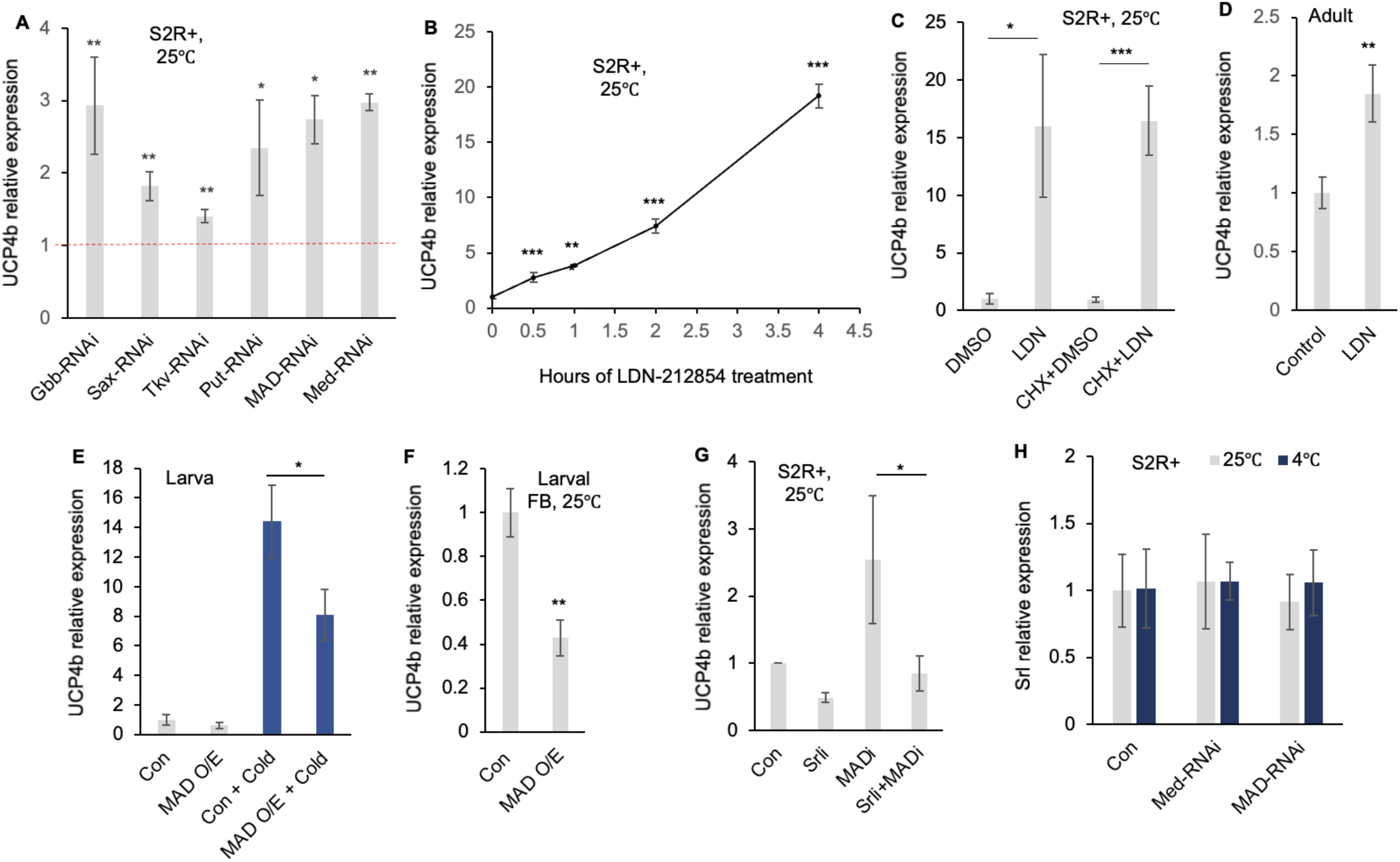
BMP signaling suppresses *UCP4b* expression. (A) *Gbb, Sax, Tkv, Put, MAD, Med* were knocked down in S2R+ cells with dsRNAs against these genes. The cells were incubated with these dsRNAs for 5 days at 25°C and then RNA was extracted and qPCR was done. Transcript levels are normalized to α-tubulin, and *UCP4b* transcript levels in these conditions are expressed relative to their levels with control dsRNA (targeting GFP) treatment. (B) *UCP4b* transcript levels in S2R+ cells are treated with 10uM LDN-212854 (at 25°C) for different durations relative to DMSO treated cells. Transcript levels are normalized to α-tubulin. (C) *UCP4b* transcript levels in S2R+ cells that were pre-treated with cycloheximide (CHX) or mock-treated for 4 hours followed by LDN-212854 treatment (at 25°C) for 4 hours. (D) *UCP4b* transcript levels in young adult males on food containing LDN-212854 compared to food containing solvent (DMSO). (E) *tubGal4, tubGal80*^*ts*^; *UAS-MAD larvae* were exposed to a permissible temperature (29°C) for 2 days to allow UAS transgene expression. Then they were exposed to cold (14°C for 4 hours). Finally, *UCP4b* expression was compared with similarly treated *tubGal4, tubGal80*^*ts*^ larvae. (F) *UAS-MAD* was expressed in the larval fat body using *R4* driver. Fat body from 3^rd^ instar larvae at 25°C was dissected and transcript levels were measured. (G) S2R+ cells were treated with *MAD* and/or *Srl* dsRNA for 5 days at 25°C and then *UCP4b* transcript level was measured by qPCR. (H) S2R+ cells were treated with *MAD* and *Med* dsRNAs for 5 days followed by cold treatment (4°C for 1 hour) and *Srl* transcripts were measured. Transcript levels are normalized to α-tubulin.

### Crosstalk between BMP signaling and cold in the regulation of UCP4b

To understand the crosstalk between BMP signaling and cold in the regulation of *UCP4b*, we explored the molecular changes in response to cold exposure as well as to the pharmacological inhibition of BMP signaling. Consistent with its role in mammalian cells, LDN212854 reduces MAD phosphorylation triggered by Tkv and Tkv^act^ overexpression in S2R+ cells, thus establishing that LDN212854 can inhibit BMP signaling in *Drosophila* (Fig 4A). Interestingly, unlike LDN212854, cold exposure did not change Tkv and Tkv^act^ overexpression-driven MAD phosphorylation indicating that BMP signaling upstream of MAD phosphorylation is not affected by cold (Fig 5A), suggesting that *UCP4b* is induced through different mechanisms by cold and LDN212854. This model is supported by the observation that simultaneous application of cold and LDN212854 synergistically induces *UCP4b* (Fig 4B). Furthermore, *MAD* knockdown also results in synergistic induction of *UCP4b* expression when combined with cold exposure (Fig 4C). Interestingly, both cold exposure and LDN212854 reduce MAD binding to the upstream regions of the *UCP4b* gene suggesting that cold exposure removes the inhibitory effect of BMP signaling on *UCP4b* by displacing MAD from the upstream regions of *UCP4b* (Fig 4D). Moreover, this displacement of MAD by cold is abrogated by *Srl* knockdown, suggesting that Srl is required for cold-mediated displacement of MAD from the upstream regions of *UCP4b* (Fig 4E). Consistent with the synergistic induction of *UCP4b* by LDN212854 and cold, flies fed a food containing LDN212854 are better protected against cold compared to the flies on control food (Fig 4F). Moreover, this protection was attenuated by *Srl* knockdown (Fig 4F). Thus, the transcriptional repression of *UCP4b* by BMP signaling under basal conditions is removed by cold acting through Srl, resulting in *UCP4b* induction.

**Figure 4.**
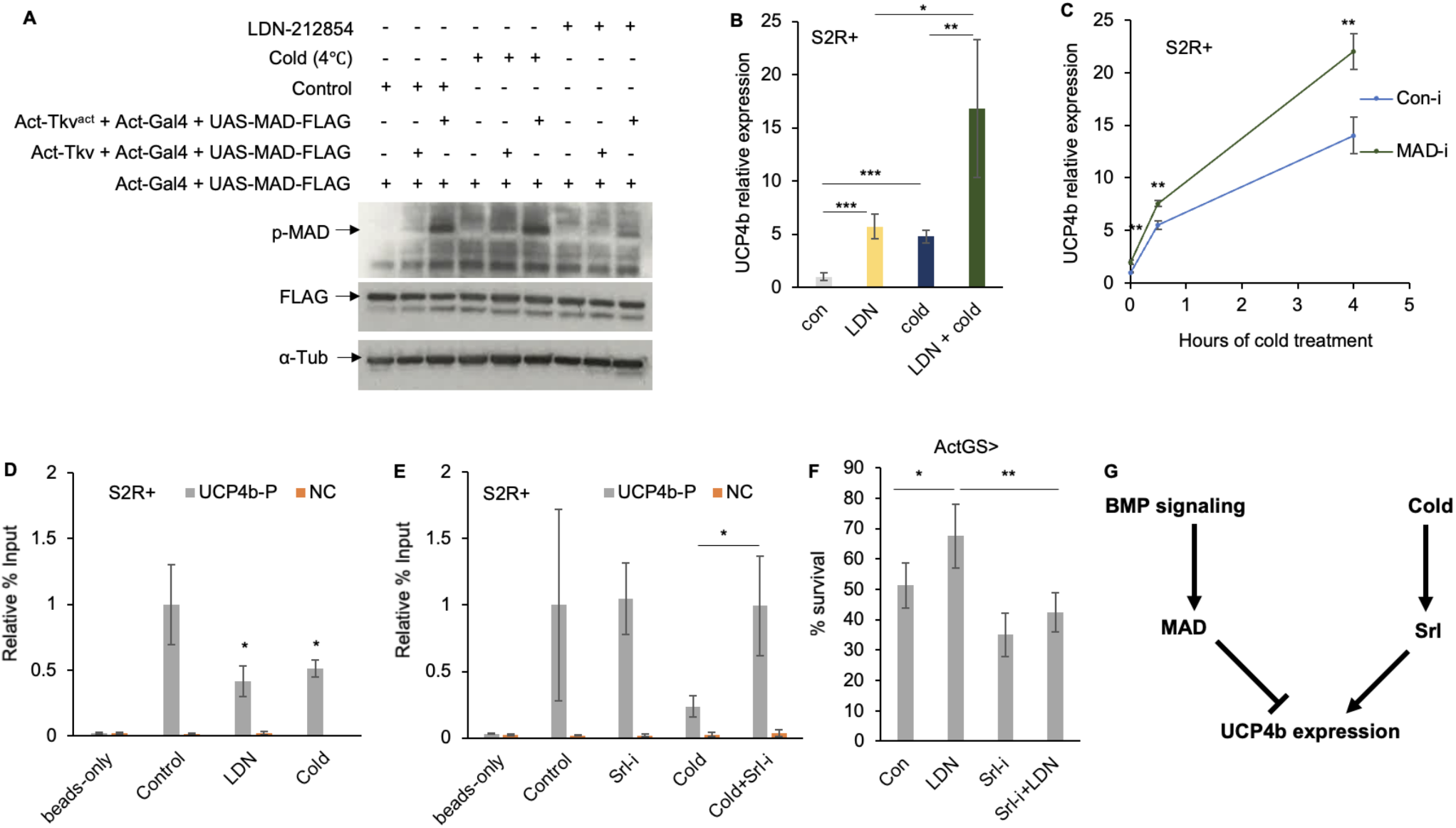
Cross-talk between cold and BMP signaling. (A) S2R+ cells are transfected with different plasmid combinations (1: Act-Gal4 + UAS-MAD-FLAG; 2: Act-Gal4 + UAS-MAD-FLAG + Act-Tkv; 3: Act-Gal4 + UAS-MAD-FLAG + Act-Tkv^act^) followed by treatment with either cold or LDN-212854. phospho-MAD, MAD-FLAG, and α-tubulin protein levels were determined using western blots. (B) *Drosophila* S2R+ cells were pre-treated with LDN-212854 (LDN) or mock-treated for 4 hours and then exposed to cold (4°C) for 2 hours. Then RNA was extracted and qPCR was performed (n=3). *UCP4b* transcript levels are normalized to α-tubulin. (C) S2R+ cells were treated with *MAD* dsRNA for 5 days followed by cold treatment (4°C) for different durations and *UCP4b* transcripts were measured. Transcript levels are normalized to α-tubulin. (D, E) Percentage enrichment for MAD binding upstream of *UCP4b* inferred by ChIP with anti-FLAG antibody. (D) MAD-binding in S2R+ cells in response to cold exposure and LDN-212854 treatment is expressed relative to MAD-binding in vehicle-treated (DMSO) cells at 25°C. (E) MAD-binding in response to cold in the presence/absence of *Srl* knockdown. (F) Cold sensitivity in adult flies after putting them on food containing LDN-212854 and/or *Srl* knockdown measured following the protocol described in Fig. 1C. (G) Model of *UCP4b* regulation by cold and BMP signaling: At normal temperature, MAD, acting downstream of BMP signaling, bind to the upstream region of *UCP4b* and suppress its transcription. Cold exposure leads to the displacement of MAD in an Srl-dependent manner resulting in *UCP4b* induction.

## Discussion

We report that, like mammalian *Ucp1, Drosophila UCP4b* is induced by cold, is suppressed by BMP signaling, and plays a role in cold tolerance and energy storage. Our studies indicate that at normal temperature, MAD, acting downstream of BMP signaling, binds to the upstream regions of the *UCP4b* gene and suppresses its transcription. Cold exposure acts through Srl to displace MAD and promote *UCP4b* transcription (Fig. 4G). *MAD* overexpression stoichiometrically counters this displacement and thereby partially blunts the cold-mediated *UCP4b* induction (as seen in Fig. 3E). On the other hand, *MAD* knockdown or LDN-mediated BMP signaling inhibition reduces this inhibitory component and thereby facilitates *UCP4b* induction (as seen in Fig. 3A and 3B). The inducing effect of cold on *UCP4b* expression synergizes with the effect of either *MAD* knockdown or LDN-mediated BMP inhibition to cause a stronger *UCP4b* induction (as seen in Fig. 4B and 4C). In addition, calcium signaling is triggered in response to cold and is important for cold-mediated *UCP4b* induction.

Recent studies have shown that non-shivering thermogenesis plays a role in cold tolerance in *Drosophila*. Mammalian *Ucp1*, which is induced in response to cold, plays a crucial role in non-shivering thermogenesis (2). Studies in *Drosophila* have shown that UCPs play a similar role in flies. *UCP4c* knockdown renders larvae sensitive to cold (15) whereas disrupted expression of *UCP4b* and *UCP4c* due to transposon insertion reduces cold tolerance in adult flies with *per* deficiency (16). Moreover, UCP-independent non-shivering thermogenesis is also shared between mammals and *Drosophila*. Sarcolipin uncouples ATP hydrolysis from Ca^2+^ transport into the endoplasmic reticulum (ER) lumen by interacting with Sarco/ER Ca^2+^ adenosine triphosphatase (SERCA) in mammalian muscles (23). Similarly, thyroid adenoma associated (THADA) uncouples ATP hydrolysis from the Ca^2+^ transport conducted by SERCA in *Drosophila*. This leads to reduced Ca^2+^ transport into the ER, and as a consequence, the energy from ATP hydrolysis is dissipated in the form of heat. Consistent with this observation, THADA-deficient flies have reduced energy production, are hyperphagic and obese, and are sensitive to cold (24). Thus, multiple mechanisms of non-shivering thermogenesis are shared between mammals and flies. Adding to these previous studies, we have shown that in addition to the evolutionarily conserved role in cold tolerance, *Drosophila UCP4b*, like mammalian *Ucp1* is also induced by cold.

The regulation of mammalian *Ucp1* by TGFβ/BMP signaling is complex. On one hand, Smad3, acting downstream of TGFβ signaling suppresses *Ucp1* expression in the white adipose tissue (9). Moreover, Smad3 binds to the promoter region of *Ppargc1a*, a transcriptional inducer of *Ucp1*, and represses *Ppargc1a* transcription in white adipocytes (9). Similarly, BMP4 signaling suppresses *Ucp1* expression in brown pre-adipocytes (10). Interestingly, BMP4 also represses the transcription of *Ppargc1a* in brown pre-adipocytes (10). On the other hand, BMP7 promotes the expression of *Ucp1* and regulators of brown fat fate including *Ppargc1a* in brown pre-adipocytes (25). Similarly, TGF-β2, an exercise-induced adipokine, promotes the expression of both in *Ucp1* and *Ppargc1a* BAT (26). Moreover, hepatic overexpression of *activin E* promotes *Ucp1* expression in inguinal WAT, mesenteric WAT and interscapular BAT (27). In contrast, hepatic overexpression of *activin E* promotes *Ppargc1a* expression only in interscapular BAT but not in inguinal WAT and mesenteric WAT (27). Thus, various TGFβ and BMP ligands have different effects on *Ucp1* expression in different adipose tissues. Moreover, the induction of *Ucp1* is mirrored by an induction of *Ppargc1a* in many cases but not all the time. Here, we report that, as in mammals BMP signaling regulates the expression of a *Drosophila* UCP (*UCP4b)*, highlighting the evolutionary conservation of this mechanism.

We have demonstrated that there are similarities between the regulation of mammalian and *Drosophila UCPs*. Similar to the mammalian *Ucp1, Drosophila UCP4b* is induced by cold. Interestingly, we found that *UCP4b* is induced by cold in a cell-intrinsic fashion, which is reminiscent of the cell-autonomous induction of *Ucp1* in mammalian white adipocytes (8). *Ppargc1a* is important for transcriptional activation of *Ucp1* in BAT in response to cold (6). Similarly, we have found that PGC1α is required for cold-mediated induction of *UCP4b*. In contrast, TGFβ/BMP signaling represses mammalian *Ucp1* expression (9, 10). Consistent with that observation, we found that BMP signaling also represses *UCP4b* expression in *Drosophila*. Thus, the antagonistic pattern of regulatory effects of cold and BMP signaling on UCPs is shared between *Drosophila* and mammals. However, we found differences between the regulation of mammalian *Ucp1* and *Drosophila UCP4b* as well. *Ppargc1a* is induced by cold-induced β-adrenergic stimulation in BAT and promotes *Ucp1* induction (6). Similarly, *Ppargc1a* is also induced by cold cell-autonomously in white adipocytes (8). However, *Drosophila Srl* was not induced by cold. Interestingly, *Ppargc1a* is suppressed by both Smad3, the effector of TGFβ signaling and BMP4 in WAT and brown pre-adipocytes, respectively (9, 10). However, we found that, although Srl is required for *UCP4b* induction under BMP signaling loss-of-function conditions, it is not induced under those conditions. These observations highlight that, while the regulatory networks controlling *UCPs* share similarities between flies and mammals, the detailed molecular mechanism has partially diverged through evolution.

It will be important to elucidate how *Drosophila* cells sense cold and trigger calcium signaling. The observation that Srl is important for cold-mediated *UCP4b* induction, but is not induced by cold, combined with the finding that cold-mediated *UCP4b* induction is not affected by cycloheximide-mediated inhibition of protein-synthesis, indicates that post-translational modification of Srl might be involved in cold-mediated *UCP4b* induction. Alternatively, post-translational modification in interacting partners of Srl could also play role in this context. Post-translational modifications of PGC1α, in addition to its cold-mediated induction, play a role in *Ucp1* induction in mammals (28). Future studies in *Drosophila* will illuminate whether similar processes are involved in cold-mediated induction of *UCP4b*. Finally, as we found that LDN212854, a BMP signaling inhibitor synergistically induces *UCP4b* when combined with cold exposure, it will be of interest to determine how these signals interact and influence *Ucp1* expression in mammalian WAT and mammalian energy homeostasis.

## Materials and method

### Fly strains, food, and cold treatment

*Drosophila* were raised in a humidity-controlled incubator at 25°C with 12/12 hr dark/light cycles on standard lab food containing 15 g yeast, 8.6 g soy flour, 63 g cornmeal, 5 g agar, 5 g malt, 74 mL corn syrup per liter. A high-sugar diet (15 w/v HSD) was prepared by adding 10 g of Sucrose to 50 g of standard lab food. LDN-food was prepared by mixing LDN-212854 with the standard lab food to a final concentration of 200uM.

Young adult flies were fed on this food for 6 days before RNA extraction and qPCR analysis. The cold sensitivity assay was carried out by exposing young adult flies in food vials (3 replicates, 20 each, sexes separated after mating) to 4°C for 6 days and then were brought back to 25°C and immediately transferred to new vials. Survivors were scored after 4 hours. The following *Drosophila* strains were obtained from the Bloomington Drosophila Stock Center (BDSC): *UCP4b-RNAi* (66999), *Srl-RNAi* (33914, 33915). The *Srl* overexpression line is described by Rera et al (29). We generated the *UCP4b* overexpression fly line by integrating a plasmid expressing *UCP4b-RA* isoform with C-terminus HA tag under UAS promoter into the attP2 site.

### RT-qPCR

mRNA was prepared from whole flies, larvae, larval tissue, and from S2R+ cells using Trizol (Invitrogen) followed by DNase (TurboDNA free kit, Invitrogen) treatment and RNA purification with “RNeasy kit” (Qiagen). cDNA was synthesized using iScript cDNA Synthesis Kit (Bio-Rad). Real-time quantitative PCR (RT-qPCR) experiments were carried out using the Biorad CFX 96/384 machine. The iQ SYBR green supermix was used following the manufacturer’s protocol. ‘Delta-delta Ct’ method was used to calculate fold change in experimental conditions. Three biological replicates were used and the transcript levels were normalized to *Drosophila* α*-tubulin* and *RpL32*. The sequences of primers used in qPCR are provided in Supplementary Dataset S1.

### Cell culture, transfection, dsRNA treatment, and drug treatment

*Drosophila* S2R+ cells were cultured in Schneider’s medium with 10% fetal bovine serum (FBS) at 25°C. S2R+ cells were transfected with plasmid DNA with Effectene transfection reagent (Qiagen) following the manufacturer’s protocol. The plasmids expressing FLAG-tagged MAD, Tkv, and activated Tkv were a generous gift from Dr. Edward Eivers (30). A *UCP4b* overexpression stable cell line was generated by integrating a plasmid expressing UCP4b-RA isoform with a C-terminus HA tag under the metallothionine promoter into S2R+ cells. Knockdown of specified genes in S2R+ cells was achieved through treatment with dsRNAs that were synthesized using the MEGAscript T7 Transcription Kit (Invitrogen) and purified with “RNeasy kit” (Qiagen). S2R+ cells were treated with dsRNAs following the ‘bathing’ method (https://fgr.hms.harvard.edu/drsc-cell-rnai). Cells were treated with LDN-212854 (Sigma), cycloheximide (Sigma), actinomycin D (Calbiochem), and BAPTA-AM (Sigma) at specified concentrations and for the indicated time points. Primer sequences used for dsRNA-mediated RNA Interference are provided in Supplementary Dataset S2.

### TAG assay

The specified number of animals or dissected tissue were homogenized in 250uL cold PBS/0.05% Triton-X (PBS-TX) with 1 mm Zirconium beads (Next Advance, ZROB10) using Bullet Blender (Next Advance). Cell debris were removed using centrifugation and TAG level was measured with Infinity Triglyceride Reagent (Thermo Fisher Scientific). 5uL of homogenate or glycerol standard (Sigma) was added to 150uL of Triglyceride Reagent for each replicate in a 96-well plate. After 10 min incubation at 37°C, the absorbance at 520 nm was recorded using a plate reader. The same homogenate was used for protein estimation using the Pierce BCA assay kit (Thermo Fisher Scientific). 2uL of homogenate or albumin standard (Thermo Fisher Scientific) was added to 200uL Pierce BCA solution for each replicate in a 96-well plate. After 30 min incubation at 37°C, the absorbance at 562 nm was recorded using a plate reader.

### TMRM assay

Transfected and copper sulfate (100uM)-or mock-treated cells were washed with PBS and plated in 96-well plates. Then cells were treated with 2uM oligomycin A (Sigma) and stained with 10nM Tetramethylrhodamine, Methyl Ester, Perchlorate (TMRM) (Thermo Fisher Scientific) in complete Schneider medium for 30 minutes at 25°C. Next, cells were washed with PBS and imaged using GE IN Cell Analyzer 6000 Cell Imaging System. 3 random fields were chosen for each of the 3 biological replicates for each condition.

### Western blot

Transfected and subsequently cold- or LDN-212854-treated S2R+ cells were directly boiled in SDS sample buffer, run on a 4%–20% polyacrylamide gel (Bio-Rad), and then transferred to an Immobilon-P polyvinylidene fluoride (PVDF) membrane (Millipore). The membrane was blocked with 5% BSA in TBST (TBS with 0.1% Tween-20) at room temperature for 1 hour and then probed with the primary antibody in 1X TBST with 5% BSA overnight. This was followed by incubation with an HRP-conjugated secondary antibody. The signal was detected by chemiluminescence (Thermo Fisher Scientific). *Drosophila* phospho-MAD was detected using an anti-Smad3 antibody (Abcam, ab52903), MAD-FLAG was detected using anti-FLAG-MAb (Sigma, F3165) whereas α-tubulin was detected using anti-alpha-tubulin MAb (Sigma, T5168).

### ChIP

Chromatin immunoprecipitation (ChIP) was performed using SimpleChIP® Plus Enzymatic Chromatin IP Kit (Cell Signaling Technology, 9005S, Magnetic Beads) following the manufacturer’s protocol. The chromatin from S2R+ cells that were transfected with Act>Gal4 and UAS-MAD-FLAG plasmids and treated in specified combinations of dsRNAs, drugs, and cold was sheared using a Bioruptor (Diagenode) at high frequency for 30 cycles of 30 s ON, 30 s OFF. The chromatin was precipitated using anti-FLAG-MAb (Sigma, F3165). The relative enrichment of the purified DNA was analyzed using qPCR with primers targeting the upstream region of the *UCP4b* gene. Primers (NC) targeting a gene desert on chromosome 2R of *Drosophila melanogaster* were used as a negative control (31). The sequences of primers used in qPCR are provided in Supplementary Dataset S1.

## Acknowledgments

We are grateful to Dr. Edward Eivers for reagents. We thank Muhammad Ahmad and Sudhir Tattikota for help with microscopy, and Christians Villalta for fly embryo injections. This work was supported by NIH grants 5P01CA120964 and 5R01DK121409 to NP, NIH grants R01DK123228 and R01 DK119117 to BMS and DRCRF Fellowship DRG-120-17 to PAD. NP is an investigator of HHMI.

## Notes

### Competing Interest Statement

The authors have declared no competing interest.

