## Supplemental Appendix for "Antagonistic regulation of *Drosophila* mitochondrial uncoupling protein *UCP4b* by cold and BMP signaling"

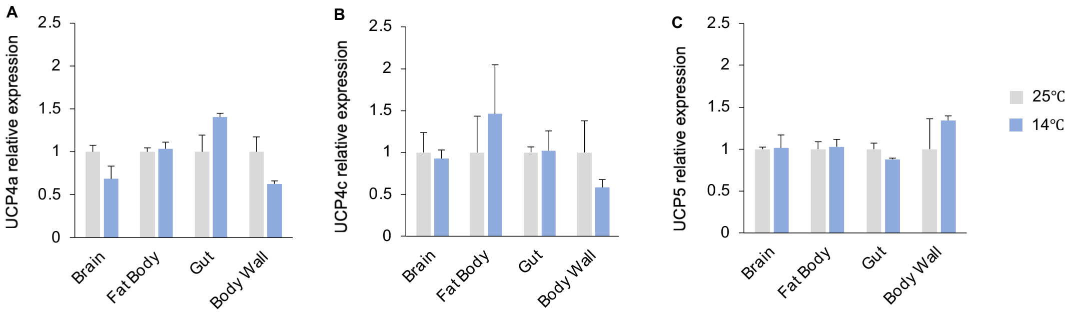


**Fig. S1. Expression of *UCP*s in *Drosophila* larval tissues in response to cold.** 3rd instar *Drosophila* larvae were exposed to cold (14°C) for 4 hours. Then the fat body, brain, gut and body wall were dissected followed by RNA extraction and qPCR.

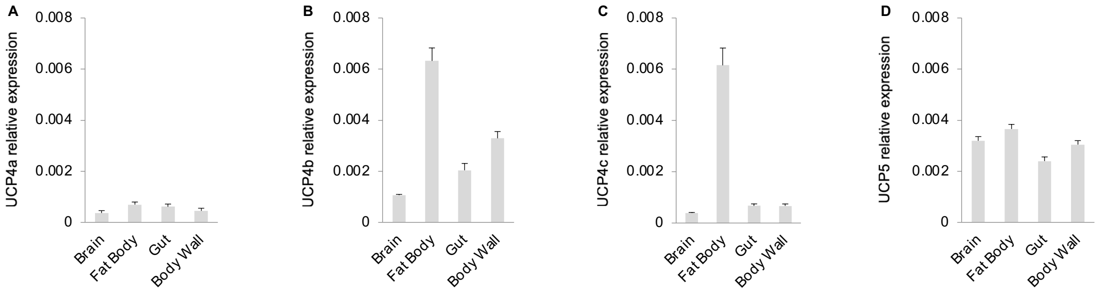


**Fig. S2. Basal-level expression of *UCP*s in *Drosophila* larval tissues.** Then the fat body, brain, gut and body wall were dissected from 3rd instar *Drosophila* larvae (at 25°C) followed by RNA extraction and qPCR.

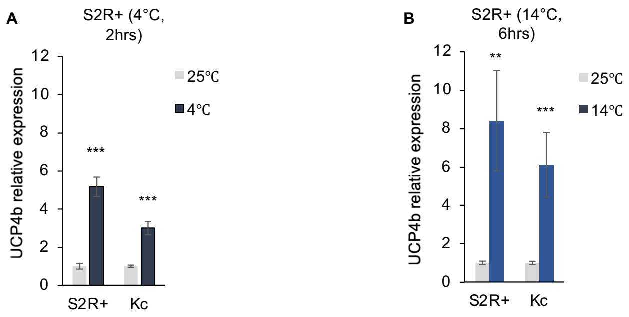


**Fig. S3**. **Cold-mediated *UCP4b* induction in *Drosophila* cultured cell lines.** Each of two different *Drosophila* cell lines (S2R+ and Kc) were exposed to different cold temperatures and for different durations (4°C for 2 hours, and 14°C for 6 hours). Then RNA was extracted and qPCR was performed.

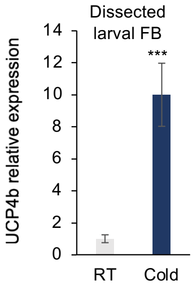


**Fig. S4**. ***ex vivo UCP4b* induction in *Drosophila* larval fat body by cold.** Dissected fat body from 3rd instar *Drosophila* larvae was incubated in cold Schneider media for 4 hours followed by RNA extraction and qPCR.

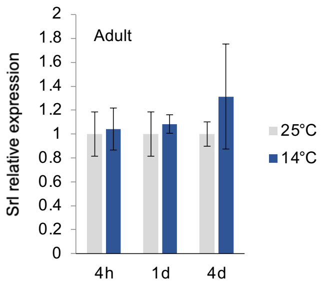


**Fig. S5**. ***Srl* is not induced in adult *Drosophila* after cold exposure.** 1 week old adult flies were exposed to cold (14°C) for different times (4 hours, 1 day and 4 days) followed by whole-body RNA extraction and qPCR.

**Movie S1.** The fat body was dissected in Schneider media at room temperature from 3rd instar *Drosophila* larvae expressing calcium-sensitive GCaMP reporter under fat body-specific FB-Gal4 driver. Then cold (4°C) Schneider media was added and the changes in calcium levels were recorded under a microscope as a movie.

**Dataset S1. Primer sequences used in qPCR**

**Dataset S2. Primer sequences used for dsRNA-mediated RNA Interference**

.
